# Neuronal excitability coordinates the timing of input/output circuit assembly

**DOI:** 10.64898/2026.07.27.740696

**Authors:** Beetsi Urrieta-Chávez, Céline Tran, Brenda García-Hernández, Samuel Domenget, Fekrije Selimi

## Abstract

How neurons coordinate the timing of migration, differentiation and synaptic integration with their specific inputs and outputs remains poorly understood. Here we show that granule cells, the brain’s most numerous neuronal population, establish immature synaptic contacts on their output targets, the cerebellar Purkinje cells, well before completing their radial migration and becoming innervated by their mossy fiber inputs. This happens at a specific stage when granule cells reach the molecular layer and begin expressing presynaptic markers and a specific NMDA receptor subtype. Chemogenetic silencing of these migrating, differentiating granule cells shows that their excitability is required for output synapse maturation. Thus neuronal activity coordinates the timing of input and output synapse development. This mechanism might underlie GRIN2B-related pathologies that include autism spectrum disorders.

---

Brain development, a protracted process that takes years in humans, relies on the coordination of multiple developmental steps to achieve the final assembly and proper functioning of complex neural circuits. It is now clear that specific genetic programs control the generation and differentiation of various neuronal populations with very diverse and specific morphological and functional properties (*1*, *2*). The neurons are generated, differentiate, migrate and settle in regions where they will be connected by input neurons. Simultaneously they grow axons that will make synaptic contacts on output targets. These events are happening in a brain that continuously changes due to its growth, in terms of cell and synapse types and numbers (*3*, *4*), but also due to the influence of the environment (*5*). Thus the development of input and output connectivity for a given neuron must be tightly regulated and coordinated so that neural circuit assembly and function occur in a timely manner.

Timing is key during brain development and might be at the core of brain evolution (*6*, *7*, *8*, *9*). Some key processes such as neurogenesis and synaptogenesis are prolonged in humans compared to other species (*6*, *10*) suggesting that changes in developmental timing contribute to increased circuit complexity. Understanding the mechanisms controlling the timing of the different steps occurring during brain development is thus a fundamental question. It also impacts our understanding of neurodevelopmental disorders. For example, the study of cell cycle durations across species has suggested that timing is a cell intrinsic property (*11*, *12*), and early increased expansion of the cortical surface area has been identified in infants who later developed autism spectrum disorders (*13*, *14*). However, the postnatal environment is key for the proper development and long-term function of brain circuits. Early childhood neglect has long-term effects on cognition and is associated with the disruption of key neural circuits (*15*, *16*). When the brain starts to respond to sensory stimuli, neuronal activity switches from purely spontaneous activity to patterns that include responses to the external world (*5*, *17*). Thus, other non-cell autonomous factors such as neuronal activity in response to changes in the external environment might play a key role in coordinating the timing of developmental processes.

Granule cells (GCs) of the cerebellum are the most numerous neuronal population in the mammalian brain, constituting more than 50% of the total number (*18*). They are a key relay for the treatment of sensorimotor information and the functions of the cerebellum, a structure now involved not only in motor adaptation and coordination learning, but also in a broad range of cognitive functions such as spatial navigation, language, and emotions (*19*, *20*, *21*, *22*). The development of GCs has been an object of intense study since Ramon y Cajal first described their differentiation (*19*, *23*, *24*, *25*). Once GC precursors (GCPs) reach the developing cerebellum, they undergo clonal expansion in the outer external granular layer (oEGL, Fig. 1A), allowing the production of a huge number of GCs. These begin differentiating in the inner EGL (iEGL), where they extend their characteristic axon, the parallel fibers (PF), while migrating tangentially (*26*, *27*). GCs then switch to radial migration and cross the molecular and Purkinje cell layers before settling in the internal granular layer (IGL) where they will receive sensorimotor inputs from the mossy fibers (*27*, *28*, *29*). GCs also make synapses on their output targets, the Purkinje cells (PCs), with each PC receiving more than 150000 PF synapses. The prevailing model has been that this step occurs once GCs have settled in the IGL (*30*). Here we aimed to revisit this view and answer the following questions: When are the first contacts made by GCs on PCs? How is the timing of all these developmental steps orchestrated during cerebellar development? We show a key role for neuronal excitability for this coordination, ensuring the timely maturation of GC synapses, first made on their target PCs before they finish their differentiation and before they are contacted by their input mossy fibers.

**Fig. 1.**
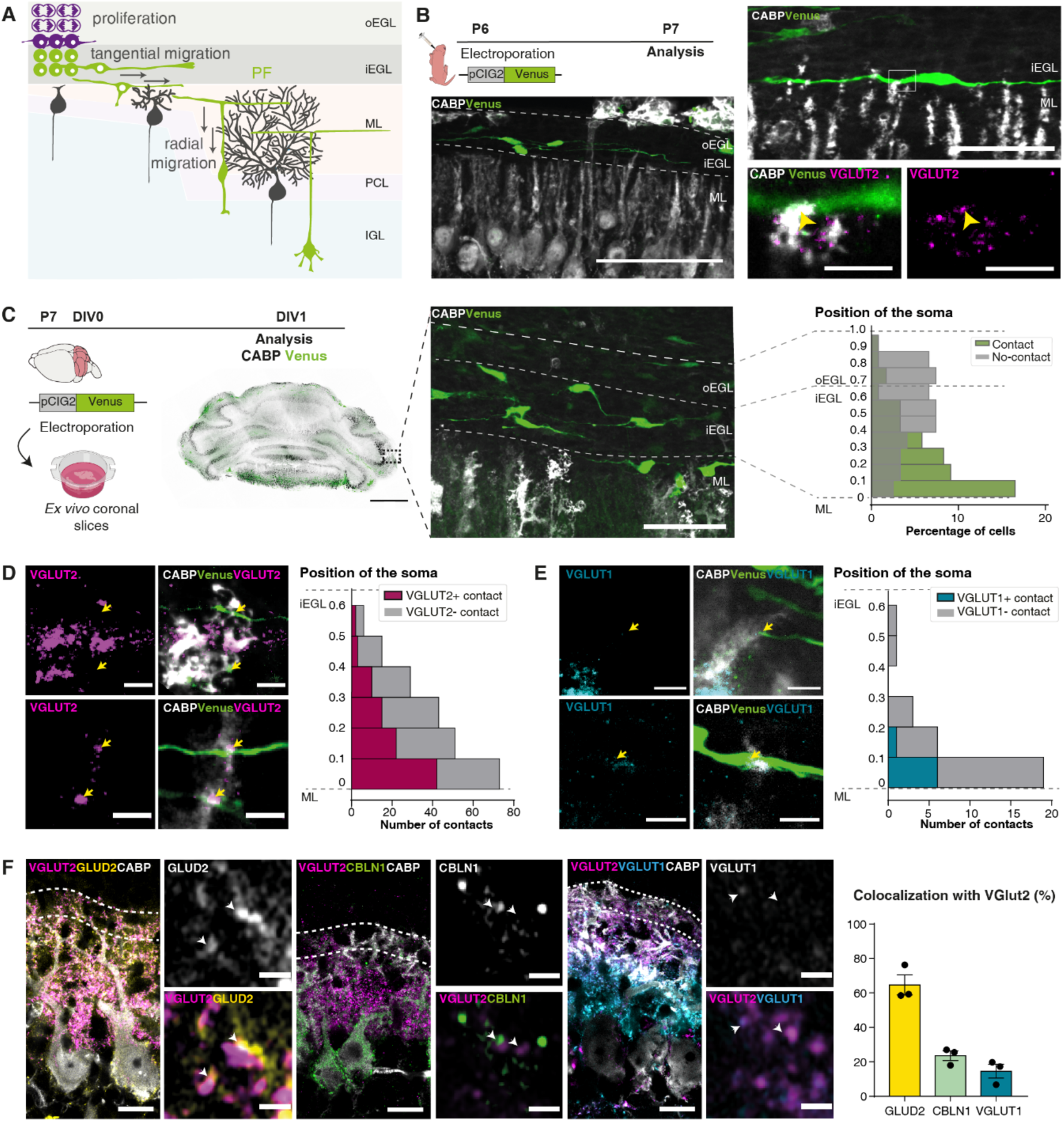
Granule cells in the external granular layer form immature synaptic contacts on Purkinje cells. **(A)** Current view of the postnatal development of the cerebellar cortex: GC progenitors (magenta) proliferate in the outer external granular layer (oEGL) and differentiate (green) in the inner external granular layer (iEGL) while migrating tangentially and extending their axons, the parallel fibers (PFs). GCs then switch to radial migration through the molecular layer (ML) and Purkinje cell layer (PCL) to reach the internal granular layer (IGL). **(B)** GC progenitors were labeled by *in vivo* electroporation of the cerebellum in P6 wild-type mice. After 24 hours, electroporated GCs expressing the fluorescent protein Venus and their newly formed PFs, PCs and their dendritic arbor and the PF immature presynaptic boutons were visualized in coronal cerebellar sections using anti-GFP, anti-CaBP and anti-VGLUT2 immunolabeling, respectively. Contacts between PCs and PFs (arrowheads) were defined by Venus and CABP colocalization in a single plane. N = 1 independent experiment. Scale bars: left = 100 µm; upper right = 50µm; lower right = 10µm. **(C-E)** *Ex vivo* electroporation of cerebella from P7 wild-type mice was followed by organotypic culture of coronal slices. At DIV1, PCs and their dendritic arbor, immature and mature PF presynaptic boutons were visualized using anti-CaBP, anti-VGLUT2 and anti-VGLUT1 immunolabeling, respectively. **(C)** Percentage of Venus-labeled GCs making contacts with PC dendrites as a function of the position of their soma relative to the ML border and normalized to the EGL thickness. The theoretical boundary between oEGL and iEGL is represented by a dashed line. Total: 121 cells, 5 mice. N = 3 independent experiments. Scale bar = 50 µm. **(D)** Number of PF/PC contacts positive or negative for VGLUT2 relative to the position of the GC soma in the EGL. Total= 230 contacts, 121 cells, 5 mice. N = 3 independent experiments. Scale bar = 10 µm. **(E)** Number of PF/PC contacts positive or negative for VGLUT1 relative to the position of the GC soma in the EGL. Total = 30 contacts, 58 cells, 4 mice. N = 1 independent experiment. Scale bar = 10 µm. **(F)** Molecular characterization of immature PF contacts located in the upper part of the developing PC dendritic arborization (top ∼12 μm, dotted line). The percentage of colocalization between VGLUT2 and other synaptic markers was quantified on parasagittal cerebellar sections of P9 wild-type mice immunolabeled for CaBP (PC somata and dendrites), GLUD2 (post-synaptic marker), CBLN1 (trans-synaptic marker) or VGLUT1 (mature PFs). N = 1 independent experiment. Scale bars: low magnification = 25µm, high magnification = 1µm.

## Results

### Pre-migratory granule cells form immature synaptic contacts on Purkinje cells

A previous study has suggested that GCs do not form PF synaptic contacts with PCs while migrating (*30*). We revisited this question by imaging individual GCs during their circuit integration. Sparse labeling of GCs with the soluble fluorescent protein Venus was performed using *in vivo* electroporation of the cerebellum in mouse pups aged 6 postnatal days (P6; Fig. 1B). After 24 hours, isolated Venus-positive GCs with growing PFs were clearly detected in the iEGL (Fig. 1B, left panel) and growing PF axons were already observed contacting PC dendrites (Fig. 1B, right panel). Quantification of the number of PF/PC contacts relative to the position of the GC soma in the EGL was then performed using *ex vivo* electroporation followed by organotypic cultures of cerebellar slices (Fig. 1C). Our results show that the number of PF/PC contacts increases as GC somata get closer to the molecular layer (ML) surface. Approximately 51% of the cells making contacts with PCs have their soma located in the 20% innermost part of the EGL (Fig. 1C). The presence of a contact by differentiating GCs on PCs is not correlated with increased length of the PF axons (Fig. S1), showing that the formation of the GC/PC contact is not simply driven by the degree of PF differentiation.

Do these contacts already have synaptic characteristics? Immature PF/PC presynaptic boutons contain the vesicular presynaptic VGLUT2 transporter while mature ones are characterized by VGLUT1 (*31*). Approximately half of the GC/PC contacts made by GCs with their soma in the lower part of the iEGL were VGLUT2 positive (Fig. 1D). As the somata of GCs get closer to the surface of the ML, the percentage of VGLUT2 positive contacts on PCs also increases, showing that the VGLUT2 presynaptic marker is progressively recruited at PF/PC contacts (Fig. 1D). VGLUT1 clusters were detected only in contacts made by GCs at the surface of the ML (Fig. 1E). Thus the initial GC/PC contacts have the characteristics of immature synapses.

What are the other molecular synaptic markers present in these immature GC/PC synaptic contacts? Co-localization of the presynaptic marker VGLUT2 with known markers of mature PF/PC synapses was quantified at the EGL/ ML border (Fig. 1F) in cerebellar sections from P9 mice. About 65% of the VGLUT2 positive boutons were associated with spines containing the postsynaptic GluD2 receptors, a key receptor involved in the formation and maintenance of PF/PC synapses (*32*, *33*). In contrast, less than 15% and 25% of the VGLUT2 positive boutons also contained the presynaptic marker VGLUT1, characteristic of mature PF/PC synapses, and the secreted synaptic protein CBLN1, a secreted synaptic organizer bridging beta-neurexin with GLUD2, respectively (*34*, *35*, *36*). Thus PF/PC synapses are born when differentiating GCs in the EGL reach the PC dendrites and form immature synaptic contacts with GluD2 containing spines. Synaptic proteins such as CBLN1 and VGLUT1 are recruited to these contacts subsequently with maturation.

### Differentiating granule cells simultaneously acquire the capacity for synaptogenesis and glutamate sensing

Classically, the EGL has been divided in two subregions with the outermost corresponding to dividing progenitors and the innermost to differentiating granule cells (*37*). In the innermost EGL, progenitors progressively generate their axons, the parallel fibers, while migrating tangentially, before switching to radial migration across PC dendrites (*25*, *26*). This raises the question as to what orchestrates the switch in the migration mode and the simultaneous initiation of synapse formation on the Purkinje cell targets.

Spatial transcriptomics followed by principal component and clustering analysis was performed using cerebellar sections at P4 (Fig. 2A), a key developmental stage characterized by intense proliferation and migration of GCPs (*25*, *38*). Focusing our analysis on segmented cells identified as part of the granule cell lineage, five different GC clusters were identified based on their differential gene expression patterns (Fig. 2B, C, D). Three subpopulations of GCs were localized in the EGL and two in the developing internal granular layer (IGL). In the IGL, one corresponded to immature GCs that have just settled and the other one to mature GCs expressing classic markers such as *Slc17a7* (sup. Data S1).

**Fig. 2.**
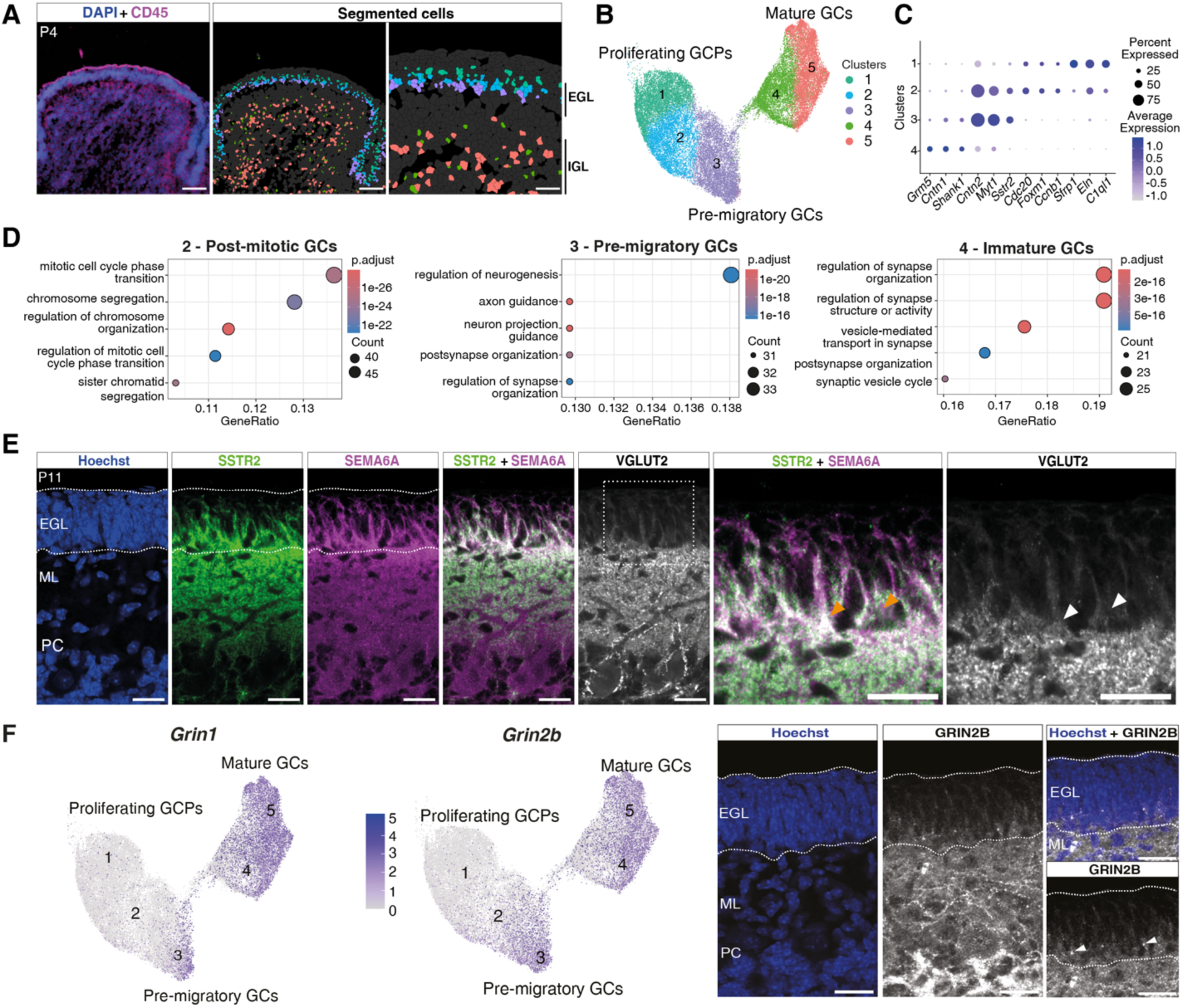
Pre-migratory granule cells acquire synapse formation and glutamate sensing capacity at the surface of the molecular layer. **(A)** Spatial transcriptomics analysis of P4 mouse coronal cerebellar sections using the Xenium In Situ platform. DAPI staining for nuclei and immunolabeling for CD45, a membrane-associated protein, were used for cell segmentation. n = 1, N = 1 independent experiment. Scale bars: low-magnification = 100µm, high-magnification (right) = 50µm. **(B)** UMAP and clustering analysis identifies five different stages of GC differentiation from proliferation of GC precursors (GCPs) to mature GCs. **(C)** Expression of key markers at each stage of differentiation of the GC lineage. Dot size indicates the fraction of cells expressing a gene, and a color scale shows the mean expression level scaled for each gene. **(D)** Dot plot representation of top enriched GO terms for biological processes in cluster 2 (post-mitotic GCs), 3 (pre-migratory GCs) and 4 (immature GCs). Dot size represents the number of annotated genes (Count) **(E)** Immunofluorescent co-labeling in parasagittal cerebellar sections from P11 mice for Hoechst (blue, nuclei), two markers of the pre-migratory GC cluster (SSTR2 in green and SEMA6A in magenta) and the VGLUT2 marker of immature PFs (white). N = 2 independent experiments. Scale bars = 20µm. **(F**) Left: Spatial transcriptomics data showing the expression of *Grin1* and *Grin2b*, genes coding two NMDA receptor subunits GluN1 and GluN2B (GRIN2B) sub-units, in the GC lineage in P4 mice. Clusters are annotated as in panel B. Right: Co-labeling for Hoechst (blue) and anti-GRIN2B (white) in a parasagittal cerebellar section from P11 wild-type mice. White arrows show GRIN2B clusters at the border of the ML. N = 2 independent experiments. Scale bars = 20µm.

In the EGL, as expected, dividing GCPs formed a layer in the outermost part and expressed markers such as *Sfrp1* showing SHH signaling responsiveness (*39*), and *Ccnb1*, a marker of proliferation (*40*). (Fig. 2C). A transition layer contained GCs that expressed both markers of mitotic cell phase transition and markers of axon differentiation and guidance such as *Cntn2*, a.k.a. TAG-1, an adhesion protein that marks growing PF axons (*41*). This transition stage likely corresponds to the previously suggested intermediate progenitor stage (*42*). Finally, the third cluster located in the inner most part of the EGL corresponds to pre-migratory GCs that have no proliferative capacity, as shown by the loss of expression of genes such as *Ccnb1* (Fig. 2C). These cells are actively differentiating with high expression of TAG-1. At this stage, differentiating GCs already start expressing regulators of synapse organization, such as *Nrxn1* and *Cbln1*, two well-known presynaptic organizers essential for synapse formation and maintenance (*34*, *35*) (Fig. S2). GO term analysis (Fig. 2D) confirmed that the pre-migratory stage is enriched in markers of differentiation in particular “axon guidance” and “regulator of synapse organization”. In addition, the observed enrichment of the GO term “postsynaptic organization” at the next stage of differentiation “immature GCs” supports the idea that mossy fibers form input synapses on GCs once these neurons settle in the IGL (*43*).

Two markers of the pre-migratory GCs in our data are SSTR2, a receptor subunit for somatostatin that decreases intracellular calcium levels in developing GCs (*44*), and SEMA6A, a guidance molecule that might be involved in the switch of GCs from tangential to radial migration (*45*). Indeed, colocalization of the SSTR2 and SEMA6A proteins peaked at the border between the EGL and the molecular layer, corresponding to the top of growing PC dendrites (Fig. 2E) in accordance with the spatial transcriptomics data. Clusters of the VGLUT2 presynaptic marker were visualized at the level of highest SSTR2/SEMA6A colocalization (Fig. 2E), confirming the acquisition of synapse forming capacity at this specific stage of GC differentiation.

Our spatial transcriptomics data show that at the pre-migratory stage GCs are also characterized by the start of the expression of several glutamate receptor subunits: the NMDA receptor subunit GluN1 (*Grin1)* and GluN2B (*Grin2b)*, and the metabotropic glutamate receptor *Grm5* (Fig. 2C and 2F). *Grin2b* is specifically expressed in differentiating GCs, and a switch to *Grin2a* occurs in mature GCs (Fig. S3 and (*46*)). Indeed, previous results have shown that differentiating GCs are responsive to NMDA (*30*). Interestingly this specific expression of the GluN2B subunit in differentiating GCs is conserved in humans (Fig. S3), suggesting a conserved function for this subunit in controlling cerebellar development.

Seminal studies from Komuro and Rakic (*30*, *47*) have shown that granule cell migration is regulated by glutamate receptor signaling, in particular via NMDA receptors. In accordance, our results suggest that pre-migratory GCs become excitable when they reach the surface of the molecular layer and switch to radial migration. Thus at this specific developmental stage, pre-migratory GCs acquire simultaneously the ability to sense glutamate and become excitable, and the capacity to form synaptic contacts with their output targets.

### Excitability in the granule cell lineage controls the timing of cerebellar development

Which other developmental processes are controlled by the capacity of differentiating GCs to become excitable at this particularly important transition?

We developed a chemogenetic approach to manipulate GC excitability specifically at different stages of their differentiation using the Neurod1-Cre mouse line that drives Cre recombinase expression in the GC lineage ((*48*) and https://www.gensat.org/CREListing.html). To enable inhibition of GC excitability, we generated Neurod1-Cre;hM4Di heterozygous mice by breeding the Neurod1-Cre mouse line with the R26-hM4Di/mCitrine mouse line driving Cre-dependent expression of the inhibitory DREADD hM4Di from the Rosa26 locus (*48*, *49*, *50*), Fig. 3A and S4). As shown by the expression of the fluorescent reporter mCitrine, by P7 most cells in the granule cell lineage of Neurod1-Cre;hM4Di heterozygotes expressed the construct, from GCPs in the oEGL to mature GCs in the IGL (Fig. S4A). Intraperitoneal acute injection of the DREADD ligand Clozapine-N-oxide (CNO) decreased the percentage of cFos positive cells in the IGL of Neurod1-Cre;hM4Di heterozygotes compared to vehicle-injected controls (Fig. S4B), confirming that our strategy enabled modulation of excitability in the GC lineage. We chose to inhibit GC excitability during cerebellar development by chronic injections of CNO between P7 and P14, a period of intense GC generation, differentiation and PF synaptogenesis (*25*, *38*, *51*, *52*). This led to a clear delay in the development of the cerebellum in Neurod1-Cre;hM4Di heterozygotes compared to littermate vehicle injected controls, as evidenced by a 15% reduction in the average surface area of vermal cerebellar sections and a 14% reduction in the thickness of the molecular layer (Fig. 3A and S4; surface area mean ± SEM: vehicle 6.12 ± 1.67 mm^2^ vs CNO 5.20 ± 5.09 mm^2^, Welch’s unpaired t-test p=0.03; molecular layer thickness mean ± SEM: vehicle 118.7 ± 2.69 µm vs CNO 101.9 ± 5.02 mm^2^, Welch’s unpaired t-test p=0.012).

**Fig. 3.**
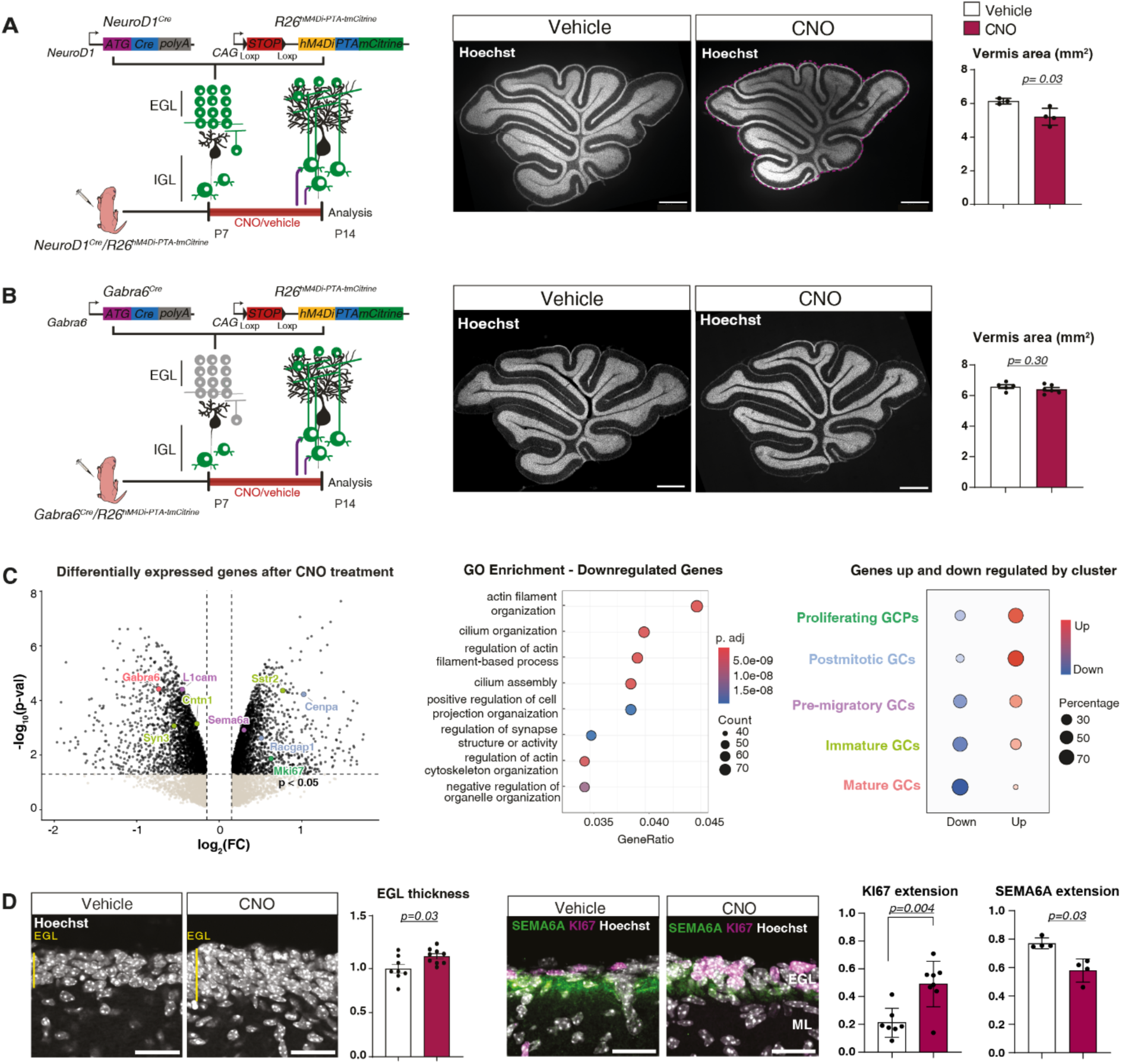
Excitability in the granule cell lineage controls the timing of differentiation. (**A**) Left: Specific expression of the inhibitory DREADD hM4Di in the GC lineage was obtained by breeding the Neurod1-Cre mouse line with a mouse line allowing the Cre-dependent expression hM4Di together with the fluorescent reporter mCitrine to obtain Neurod1-Cre;hM4Di heterozygotes. The DREADD ligand, Clozapine-N-oxide (CNO), was administered twice daily from P7 to P14. Right: The mean total surface area of parasagittal cerebellar sections taken in the vermis of P14 mice was assessed (as shown by the dotted purple line in the CNO image) after Hoechst staining. N = 1 independent experiment. Scale bars = 500μm. (**B**) Same analysis as in A but on mice expressing the inhibitory DREADD specifically in mature GCs (obtained using the Gabra6-Cre mouse line instead of the Neurod1-Cre line). N = 2 independent experiments. Scale bars = 500μm. **(C)** Left: Volcano plot representation of gene expression changes detected by bulk RNA sequencing of cerebellum extracts from Neurod1-Cre;hM4Di mice at P14 (CNO versus vehicle-treated). Vertical lines correspond to a ±10% relative expression change (log₂FC ≈ ±0.14). Some genes highly expressed in the different clusters of differentiating GCs are highlighted; Middle: GO term enrichment analysis of the differentially expressed genes (DEGs) classified according to the term “Biological Processes”. Right: Dot plot representing the percentage of genes up or down-regulated amongst the top markers characterizing each GC cluster (Same color coding as in the volcano plot). **(D)** Left: EGL thickness (normalized to vehicle-treated) measured in Hoechst-stained parasagittal cerebellar sections of P14 vehicle (n = 8) or CNO (n = 9) treated mice. N = 3 independent experiments. Right: Relative extension of the proliferating layer (KI-67 staining relative to the EGL thickness) and pre-migratory layer (SEMA6A staining relative to the EGL thickness) measured on immunolabeled parasagittal cerebellar sections of P14 mice. N = 2 independent experiments. Scale bars = 20µm.

To test whether this delayed development could result from decreased excitability in differentiated mature GCs, we generated another mouse model using the Gabra6-Cre mouse line as a driver for hM4Di specific expression in mature GCs in the IGL (*53*) (Fig. 3B and S5). No changes in the development of the cerebellum were detected after a one-week chronic reduction of GC excitability in this model, neither in the surface area of the vermis (Fig. 3B; mean± SEM: vehicle=6.588 ± 0.119 mm^2^, n = 5 vs CNO: 6.407 ± 0.115 mm^2^, n = 6, Welch’s unpaired t-test, p = 0.30) or the thickness of the molecular layer (Fig. S5). Thus decreased excitability in differentiating GCs, but not in mature ones, has profound effects on the cytoarchitectural development of the cerebellum.

The consequences of decreased excitability in differentiating GCs were further characterized by transcriptomic analysis using bulk RNA sequencing of cerebellar extracts from P14 Neurod1-Cre;hM4Di heterozygotes treated with CNO compared to vehicle-treated littermates (Data S2, Fig. 3C, left). Differential gene expression analysis (Fig. 3C, left) followed by global GO term analysis (Fig. 3C, middle) showed 435 downregulated genes in the cerebellum of CNO-treated animals with an enrichment for GO terms related to “actin filament organization”, “cilium organization” (*Sstr3*, *Ift80*, *Cep290*) and “regulation of synapse structure or activity” (*Gabra6*, *L1cam*). This results is in agreement with our observation of delayed maturation of the cerebellum in CNO-treated Neurod1-Cre;hM4Di mice. Conversely, multiple nuclear and cell cycle-related genes, including *Cdk1*, *Cdc14a*, and *Mki67* were upregulated after a one-week chronic inhibition of excitability in differentiating GCs (Data S2).

To determine at which specific stage decreased excitability affected GC differentiation, we cross-referenced the differentially expressed genes with cluster markers assigned by our spatial transcriptomics analysis. This analysis revealed that a majority of the markers of proliferating GCPs (70.6%) and intermediate (81.3%) progenitors were upregulated, while markers of immature (66.7%) and mature (87.5%) GCs were mostly downregulated (Fig. 3C, right). In accordance, the thickness of the EGL in lobule IX of the cerebellum was significantly increased by 14% at P14 in CNO-treated NeuroD1Cre/hM4Di heterozygous mice, compared to vehicle-treated controls (Fig. 3D; mean± SEM: vehicle= 23.31 ± 3.25, n = 8 vs CNO=26.6 ± 2.01, n = 9; Welch’s unpaired t-test p = 0.03). No significant increase in the EGL thickness was observed at P11 (Fig. S4), suggesting that increased EGL thickness at P14 is a result of a progressive slowdown of GC development, rather than a blockade at a specific stage. This increase in the size of the EGL at P14 was accompanied by a 19.4 % reduction in the relative thickness of the SEMA6A layer and a 93.10% increase in the relative thickness of KI67 labeling, a marker of proliferating GCPs, confirming reduced EGL maturation (Fig. 3D; normalized SEMA6A extension mean ± SEM: vehicle=0.754 ± 0.038, n = 4, vs CNO=0.604 ± 0.081, n = 4, Mann-Whitney test p=0.02; normalized KI-67 extension mean : vehicle=0.299 ± 0.102, n=7, vs CNO=0.567 ± 0.106, n=7, Welch’s unpaired t-test p=0.0004).

Finally, gene markers of the pre-migratory stage of GC differentiation were equally up- and down regulated (Fig. 3C, 54.5% versus 45.5% respectively). Thus chronic inhibition of GCP excitability delays cerebellar development by specifically slowing down the maturation of pre-migratory GCs at the moment when they switch to radial migration.

### Excitability in differentiating granule cells controls the maturation of output connectivity

GCs connect with their input mossy fibers when they reach the internal granular layer (*29*). Since pre-migratory GCs acquire the ability to form immature synaptic contacts with their targets simultaneously with the ability to respond to glutamate, a compelling hypothesis is that glutamate sensing coordinates the timing of GC migration, and thus of input connectivity, with the maturation of their output connectivity.

The maturation of PF/PC synapses can be monitored as the presynaptic vesicular glutamate transporter in PF boutons switches from VGLUT2 to VGLUT1 (*31*). At P14 in control heterozygous mice, the maturation of PF/PC synapses is visible as an inside-out decreasing gradient of VGLUT1 immunolabeling in the ML, while VGLUT2 positive PF boutons are mostly located below the remaining EGL in the upper 20% part of the ML (Fig. 4).

**Fig. 4.**
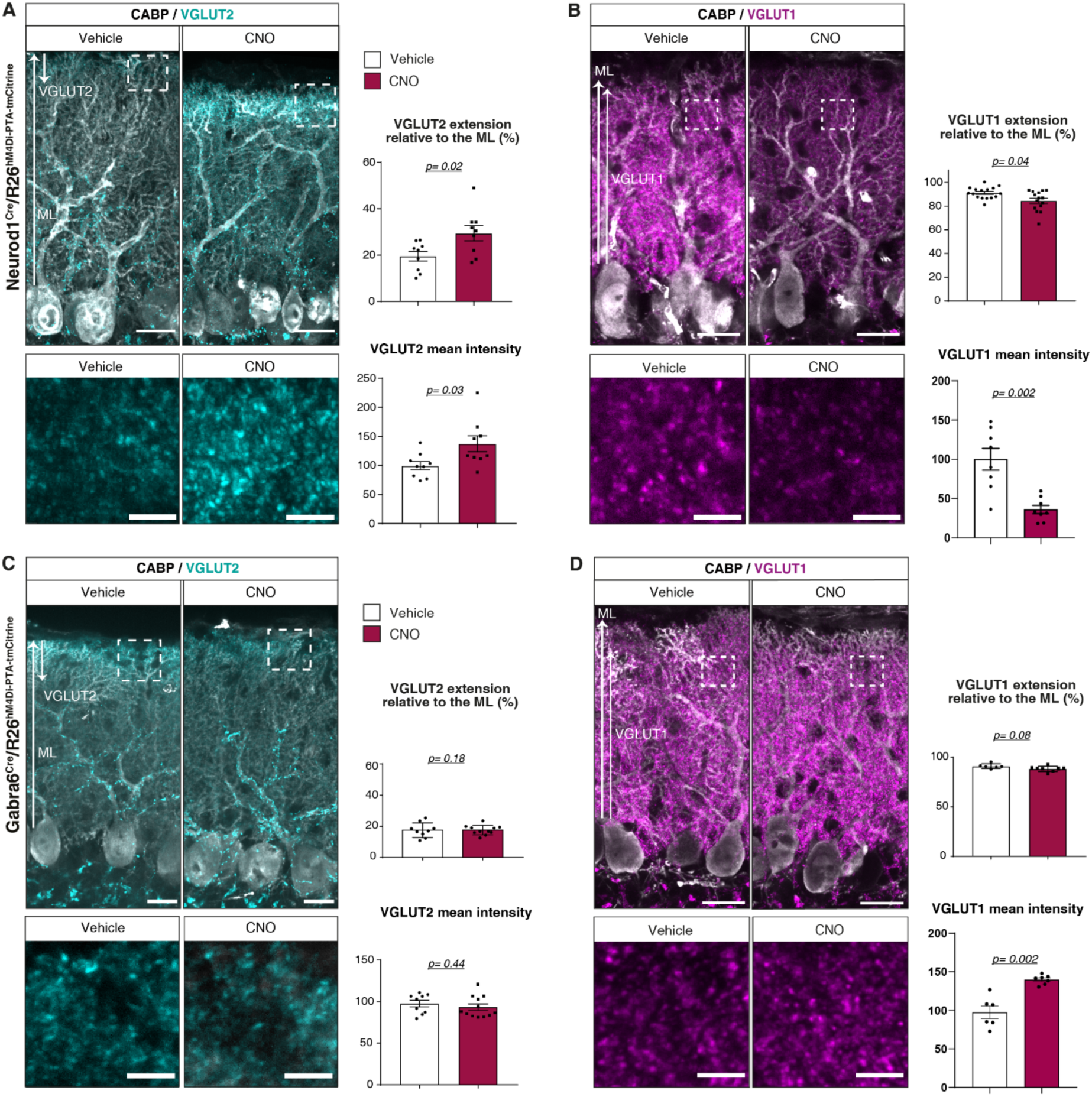
Excitability in pre-migratory granule cells controls the timing of output connectivity maturation. **(A-B)** PC somata and dendrites, and associated immature and mature PF presynaptic boutons were identified using anti-CaBP (gray), anti-VGLUT2 (cyan) and anti-VGLUT1 (magenta) immunostaining respectively in parasagittal cerebellar sections (vermis, lobule IX) of P14 Neurod1-Cre;hM4Di mice chronically treated with CNO or vehicle. The extension of the VGLUT2 or VGLUT1 PF territory relative to the total ML height was quantified as shown by the arrows. The mean fluorescence intensity of VGLUT1 and VGLUT2 clusters were quantified in the upper ML (PF-only territory, ROI, rectangle) and expressed relative to the mean of vehicle group. VGLUT2: vehicle n = 9, CNO n = 9; N = 4 independent experiment.; VGLUT1:vehicle n = 8, CNO n = 8; N = 3-5 independent experiment. Scale bars = 25μm, high-magnifications = 5μm. **(C-D)** Same analysis using vermal parasagittal cerebellar sections of P14 Gabra6-Cre;hM4Di mice. VGLUT2 extension: vehicle n = 9, CNO n = 11; N = 5 independent experiments;;VGLUT2 mean intensity: vehicle: n = 9, CNO: n = 12; N = 4 independent experiments; VGLUT1 extension: vehicle n = 6, CNO n = 8; N = 2 independent experiment.; VGLUT1 mean intensity: vehicle n = 6, CNO n = 7; N = 2 independent experiments. Scale bars = 25μm, high-magnifications = 5μm.

After one-week chronic inhibition of excitability, a clear impairment in PF/PC maturation was evident at P14 in Neurod1-Cre;hM4Di heterozygous mice, with a 50% increased extension of the VGLUT2 PF territory relative to the ML thickness (Fig. 4A; mean ± SEM vehicle: 19.5 ± 2.0 µm vs CNO: 29.44 ± 3.26 µm, Welch’s unpaired t-test p = 0.02). Conversely, the extension of the territory of VGLUT1 positive PFs was decreased by about 7% (Fig. 4B; mean ± SEM, vehicle: 91.19 ± 1.22, vs CNO: 84.59 ± 2.17 µm, Welch’s unpaired t-test p = 0.04). In addition, the mean VGLUT1 intensity in the ML was decreased by 37.5% (Fig. 4B, mean intensity normalized to vehicle ± SEM, vehicle: 100 ± 13.90, n = 8 vs CNO: 35.91 ± 5.30, n = 8, Welch’s unpaired t-test p = 0.002), and the mean VGLUT2 intensity increased by 37% (Fig. 4A, mean ± SEM, vehicle: 100 ± 7.02, vs CNO: 137.4 ± 13.57, Welch’s unpaired t-test p = 0.03), in agreement with delayed maturation of PF/PC synapses. Interestingly the expression of key presynaptic organizers, *Nrxn1* and *Rtn4r* (*54*), but not of *Cbln1*, was significantly reduced in cerebellar extracts from CNO-treated NeuroD1Cre;hM4Di heterozygous mice compared to the vehicle-treated controls (Data S2). Thus excitability in differentiating GCs controls PF/PC synaptic maturation by regulating the expression of synaptic proteins both at the transcriptional and post-transcriptional levels.

To test whether decreasing excitability in mature GCs has the same effect on PF/PC synapse maturation, we performed the same analysis using Gabra6-Cre;hM4Di heterozygotes. After one-week of chronic treatment with CNO, no significant changes were detected when compared to vehicle-treated controls, neither in the extension of the VGLUT1 and VGLUT2 PF territories relative to the ML height, nor in the mean VGLUT2 intensity ((Fig. 4C and 4D). The VGLUT1 mean intensity was increased by 43.5%, suggesting a homeostatic compensation for decreased excitability of mature GCs. Thus, comparison of the results obtained with the two chemogenetic models shows that excitability in differentiating GCs, not mature GCs, simultaneously controls the timing of GC differentiation at the pre-migratory stage and the timing of maturation of their output connectivity on target PCs.

## Discussion

Studies of the many steps of neural circuit development, from proliferation and differentiation of neurons to synaptogenesis, have revealed the underlying molecular mechanisms for each step individually and their contribution to neurodevelopmental diseases (*1*, *2*, *4*, *23*, *55*, *56*). However, how these steps are orchestrated from the viewpoint of an individual neuron in a developing brain remains unknown. Our work reveals that for granule cells (GCs) in the developing cerebellum, these steps are coordinated by neuronal excitability. Differentiating GCs start making output synapses on PCs when they are still at the pre-migratory stage. This is also when they acquire the ability to sense glutamate and when neuronal excitability coordinates the pace of differentiation, switch to radial migration and the maturation of their output synapses.

Mossy fibers, the inputs of GCs, originate from several brainstem nuclei, and reach the cerebellum before birth (*43*). A combination of repulsive and attractive molecular signaling prevents them from entering the EGL and ML (*29*, *43, 57*), and controls their specific connectivity on GCs. For example, expression of CDH7 prevents mossy fiber overgrowth in the ML and promotes synaptogenesis with GCs (*58*). Indeed, *Cdh7* is expressed by mature GCs in the IGL (Data S1), confirming that input connectivity on GCs occurs when they have completed their migration and differentiation. The timing of input connectivity is thus tightly regulated by both intrinsic and extrinsic factors. Contrary to the previous view (*30*), our results show that GCs make immature synapses on PCs while they have not completed their differentiation and are still far from their final destination. These output synapses must then mature in coordination with the timing of migration and differentiation of GCs, to ensure the proper and timely transfer of information from sensorimotor inputs via mossy fibers to the output of the cerebellar cortex. Indeed, neural circuit activity during this period is changing in response to environmental stimuli and circuit maturation itself (*51*), leading to circuit adaptation and proper cognition and function.

Our work shows that neuronal excitability is key for the coordination of input/output connectivity development. This adds to the other roles that spontaneous and evoked neuronal activity have been shown to play in shaping sensory circuits (*5*, *17*). Neuronal excitability can regulate GCs development in a stage-dependent manner. In immature GCPs, membrane depolarization by KCl or by GABA promotes proliferation through Ca^2+^ increase. At late postmitotic stages NMDA receptor activation regulates axonal growth, migration and survival (*30*, *59*, *60*). Our in vivo work shows that neuronal excitability is key for the timing of differentiation and the exit from the EGL of GCs and the timely maturation of their output synapses. A recent study has shown that postnatal neuronal activity is also key for the maturation of parvalbumin interneurons in the cortex, via the control of PGC1α-mediated transcriptional programs (*61*). NMDA receptors also control the switch from slow multipolar migration of neurons to radial glia guided migration in the neocortex (*62*). Thus the coordination of input/output connectivity maturation by neuronal activity is probably a general function throughout brain circuits.

In this regard, a key question will be to determine the neuron-specific molecular factors underlying this coordination. In GCs, different genetic programs have been shown to underlie their stepwise differentiation, involving Tet epigenetic factors (*63*, *64*). In particular bivalent epigenetic marks are abundant at the beginning of differentiation and H3K27 methyltransferases EZH1 and EZH2 are essential to regulate the timing of expression of many genes involved in axon guidance, synaptogenesis and terminal differentiation (*65*). This suggests that these marks must be controlled by activity-dependent epigenetic factors that remain to be determined and that will be key for coordinating the timing of brain circuit development.

Lack of sensorimotor stimuli during postnatal brain development leads to long-term deficits in cognition in humans (*66*). Mutations in *GRIN2B*, a key subunit for glutamate sensing in differentiating GCs, are associated with various neurodevelopmental disorders characterized by mild to profound developmental delay and intellectual deficiency, as well as epilepsy and autism spectrum disorders (*67*). Key perspectives are thus to understand to what extent these symptoms are associated with defects in the timing of connectivity maturation and the specific contribution of cerebellar circuit developmental defects.

## Acknowledgements

We would like to thank Pr. C.A. Mason and Dr. S. Sigoillot for their advice on the manuscript. We would like to thank the CIRB facilities in particular T. Piolot, J. Dumont, T. Caille, and other members of the ORION microscopy facility, A. Robert and other members of the animal facilities and F. Ladouce and other members of the administrative department. The spatial transcriptomic experiment using the Xenium technology was performed with the PHENOMIN - iCS facility, Illkirch, France. We would like to thank Antoine Valera for contributing to the quality control of probes in the spatial transcriptomics approach. Bulk RNA-sequencing was performed at the Institut Curie, PSL University, ICGex Next-Generation Sequencing Platform. We would like to thank Maxime Corbe, Sylvain Baulande and Nicolas Servant from the Bioinformatics facility of the Institut Curie for their help and advice on data analysis for the bulk RNAseq and spatial transcriptomics.

This work was supported by grants from Agence Nationale de la Recherche ANR-22-CE16-0028, Ligue Nationale Contre le Cancer 294983, Fondation pour la Recherche Médicale EQU202503020071, ECO202406019167, FDT202304016886, ANR-10-LABX-54 MEMO LIFE.

## Materials and Methods

### Animals

All animal protocols were approved by the Comité Régional d’Ethique en Expérimentation Animale (no. CEEA 59), and animals were housed in authorized facilities of the Center for Interdisciplinary Research in Biology (D750512).

Wild-type mice used in the experiments were C57BL6/J obtained from Charles River Laboratories (Wilmington, USA). The Neurod1-Cre mouse line (Tg(Neurod1-cre)RZ24Gsat/Mmucd) was a kind gift from Pr. Mary Beth Hatten (The Rockefeller University, USA). Transgene genotyping was performed using the following primers: 5’TAG GAT TAG GGA GAG GGA GCT GAA 3’ and 5’ CGG CAA ACG GAC AGA AGC ATT 3’. The Gabra6-Cre mouse line (Gabra6tm2(cre)Wwis, (*53*)) was obtained from Dr. David DiGregorio (Institut Pasteur, France). Transgene genotyping was performed using the following primers: 5’-GAT CTC CGG TAT TGA AAC TCC AGC −3’ and 5’-GCT AAA CAT GCT TCA TCG TCGG −3’. The hM4Di DREADD mouse line, B6.129-Gt(ROSA)26Sortm1(CAG-CHRM4*,-mCitrine)Ute/J, was obtained from the Jackson laboratory (Bar Harbor, USA). Genotyping was performed using the following primers: 5’CGA AGT TAT TAG GTC CCT CGA C 3’ and 5’ TCA TAG CGA TTG TGG GAT GA 3’. Neurod1-Cre or Gabra6-Cre homozygous mice were crossed with hM4Di DREADD homozygous mice in order to obtain Neurod1-Cre;hM4Di or Gabra6-Cre;hM4Di double heterozygous mice. Mice from both sexes were used in our experiments.

Clozapine-N-oxide (3µg/g) or vehicle (saline solution NaCl 0.9%) was injected intraperitoneally from P7 to P14 twice per day.

Harmaline (50mg/kg) dissolved in physiological serum was injected intraperitoneally 1h30 minutes before perfusion.

### Ex vivo Cerebellar Electroporation

P7-P8 cerebella were dissected from C57BL/6J mouse brains in Hanks balanced salt solution (HBSS; ThermoFisher #14170112) containing 2.5mM HEPES (pH 7.4), 30 mM D-glucose, 4 mM NaHCO3, added with 1 mM CaCl2, 1 mM MgSO4 on ice. The cerebella were incubated with the pCIG2-Venus plasmide DNA diluted at 0.5 μg/μL in HBSS supplemented with D-glucose (final concentration: 46 mM) for 15–20 min on ice. Electroporation was performed in the electroporation chamber (Protech International, Inc. CUY520P5 platinum electrode L8xW5xH3 with 5-mm gap) using an NEPA21 type II electroporator with the following parameters: 50 msec at 80 V for a total of five pulses with an interval of 500 msec between pulses and a poring pulse of 80V / 30 msec pulse length/ 450msec pulse interval/ 1 pulse/ 10% decay rate/ polarity +/-. Following electroporation, the cerebella were recovered in HBSS supplemented with D-glucose for 10 min on ice. Coronal slices of 200 µm were cut using a Campden Ci 7000 smz microtome in HBSS, and placed on Millicell culture plate inserts (CM 0.4-μm) for culture in the following medium: Basal Medium Eagle (ThermoFisher #21010046), 25mM D-glucose, 1× GlutaMAX (ThermoFisher #35050061), 1x Insulin-Transferrine-Selenium (ThermoFisher #41400045), 1X Penicillin-Streptomycin (ThermoFisher #15140148). Slices were cultured for 24-48 hours at 37°C and 5% CO2. Slices were fixed with 4% paraformaldehyde in phosphate-buffered saline for 20 minutes at room temperature before being processed for immunostaining.

### In vivo electroporation

P6 C57BL/6J mice were anesthetized by hypothermia on ice. 1 µL of pCIG2-VENUS plasmid (8,5 µg/µL) was injected using a calibrated pulled glass capillary in the cerebellum. Injection coordinates were defined as x= - 4.2 mm, y = −1.12 mm, z = - 2 mm. The plasmid solution was delivered slowly to minimize tissue damage. Electroporation was performed using 5 mm Tweezer-type electrodes (CUY650P5, NEPA GENE) connected to a square-wave pulse generator (NEPA21 Type II). Electrodes were positioned over the head to direct the electric pulse to the cerebellum. 7 pulses of 80 V and 50 ms duration were delivered at 150 ms intervals. The wound was closed with surgical glue and disinfected with Betadine 10%. Pups were allowed to recover on a heating pad at 37 °C before being returned to the dam.

### Antibodies

The following primary antibodies were used: mouse monoclonal anti-CaBP (1:7000; Swant, Switzerland, #300), rabbit polyclonal anti-CaBP (1:1000; Swant, #CB38), guinea pig polyclonal anti VGLUT1 (1:5000; Millipore, Massachusetts, USA, #AB5905), guinea pig polyclonal anti-VGLUT2 (1:5000; Millipore, #AB2251), mouse monoclonal anti-VGLUT2 (1:500 SynapticSystems #135421), chicken polyclonal anti-GFP (1:1000; Abcam, #ab13970), goat polyclonal anti-SEMA6A (1:300; R&D Systems #AF1615), rabbit polyclonal anti-KI67 (1:2000 Abcam #15580), rabbit monoclonal anti-C-FOS (1: 2000, SynapticSystems #226003), mouse monoclonal anti-NR2B (1:250, BD Bioscience #610416), rabbit polyclonal anti-CBLN1 (1:200, Nittobo Medical #MSFR100650). The following secondary antibodies were used: goat anti-guinea pig Alexa Fluor 568 (2μg/mL; Invitrogen, California, USA, #A11076), goat anti-guinea pig Alexa Fluor 647 (2μg/mL; Invitrogen, #A21450) donkey anti-mouse Alexa Fluor 568 (2μg/mL; Invitrogen, #A10037), donkey polyclonal anti-Rabbit Alexa Fluor 488 (2μg/mL; Invitrogen, #A21206), donkey polyclonal anti-Rabbit Alexa Fluor 647 (2μg/mL; Invitrogen, #A31573), donkey polyclonal anti-Rabbit Alexa Fluor 568 (2μg/mL; Invitrogen, #A10042), goat anti-chicken (2μg/mL; Invitrogen, #A11039).

### Immunofluorescent labelings

Brains were extracted after intracardiac perfusion of mice with 4% paraformaldehyde (PFA) in phosphate-buffered saline (PBS) solution (for anti-CaBP, anti-GFP, anti-VGLUT1, anti-VGLUT2, anti-C-FOS, anti-KI67 and anti-SEMA6A stainings) or with 9% glyoxal (from a 40% stock solution, Sigma-Aldrich #128465) and 8% acetic acid (Supelco #160250) in water (for anti-NR2B staining). Brains were post-fixed in 4% PFA/PBS or 9% glyoxal/8% acetic acid for one hour (P7) or overnight (P11, P14) at 4°C, and then cryoprotected for 48 h in a 30% sucrose/PBS solution at 4°C. After sectioning with a freezing microtome, 30μm-thick sections were washed and blocked with 4% normal donkey serum in PBS for 1 hour at room temperature. For CBLN1 immunolabeling, an antigen retrieval step was added before blocking: sections were treated using pepsin 1mg/mL (Thermo Scientific Chemicals # 3765723) /HCl 12N (Sigma-Aldrich # 258148) at 37°C for 10 min, and washed 3 times for 5 min with PBS. The primary antibodies were diluted in PBS/1% Triton X-100 (for VGLUT2 and VGLUT1 detection) or 0.4% Triton X-100, 1% donkey serum, and incubated overnight at 4°C under agitation. The sections were then washed three times for five minutes in PBS/ Triton X-100 and incubated for two hours at room temperature in the secondary antibody, diluted in PBS 1X, 1% Triton X-100, 1% donkey serum. After 3 washes in PBS/ Triton X-100, the sections were incubated for 15 minutes at room temperature with the nuclear marker Hoechst 33342 (0,2 mg/mL, Sigma-Aldrich #H6024) in PBS/ Triton X-100. After a final wash in PBS, the sections were mounted with ProLong Gold Antifade Reagent (Invitrogen, #P36930).

### Image acquisition and quantifications

Images for global cerebellar morphology were acquired using a Zeiss Axiozoom. Otherwise, image were acquired using a spinning-disk confocal CSU-W1 Zeiss microscope with a 25X, 40X, 63X or 100x objectives, using Metamorph Premier 7.8 software.

For quantification of contacts according to cell position, image stacks spanning the entire region of interest were acquired with a Zeiss CSU-W1 spinning-disk confocal microscope using a 40x objective (NA 1.4), with a pixel size of 0.16 µm (2048 x 2048 pixels, 16-bit) and a z-step of 0.3 µm, covering almost the full depth of the tissue (up to 63 planes per stack). Depending on the sample, the region of interest was either imaged as a single mosaic acquisition or captured as separate adjacent image stacks that were manually reconstructed into a composite. Electroporated cells were identified by Venus expression and manually labeled. Only cells whose entire axon length could be followed throughout the stack were included in the analysis. Because electroporated cells migrate tangentially along the external granule layer (EGL), the position of each cell was determined by tracing a line perpendicular to the soma, from the border of the soma to the nearest CaBP-positive signal marking the molecular layer (ML) border; this distance was used to estimate the soma’s position relative to the ML. Contacts were defined as the colocalization, within a single Z plane, of Venus, and CaBP (calbindin) signals with a minimum overlap of 4 pixels in XY required between signals. IN addition the signal for synaptic markers (VGluT1 or VGluT2) was also quantified in each contact. Colocalization analysis was performed in ImageJ/Fiji. Contact quantification was carried out blind to condition and independently verified by two observers.

The axon length of electroporated granule cells were measured with NeuronJ plugin, an ImageJ plugin for 2D tracing and analysis of elongated structures, allowing to follow axonal length. Briefly, PF-axons were semi-automatically traced from the soma to the axon growth cone. Trace segments were verified manually and corrected where necessary to ensure accurate path selection. Axonal length was then calculated and converted to microns.

GLUD2, CBLN1 and VGLUT1 (Fig1.E) were taken using a AiryScan 2 LSM 980 Confocal with 63X (NA: 1.4) objective. Quantitative analysis of images was performed using custom-made plugins using ImageJ Fiji open source software (https://github.com/orion-cirb/Cfos_2D_Syn_Dendrites). In brief, the background from each channel was removed, a threshold was applied and the image converted to a mask. Colocalization of the different markers with VGLUT2 was then quantified using ImageCalculator using a mask-based approach. Manual quantifications were done in blind condition.

For VGLUT1 and VGLUT2 from quantification in parallel fibers, the background of the whole image was subtracted using the Fiji built-in subtract background plugin. The signal measured was the raw integrated density (the sum of the values of the pixels in the selection) divided by the area in μm², thus providing a measure of the mean intensity per μm². The extension of the VGluT1+ or VGluT2+ parallel fiber PF territory was measured according to a defined threshold of gray values. As the signal lacked a precise boundary and faded progressively, a threshold was set in Fiji at the value corresponding to signal onset and applied consistently across all images within the same batch to determine PF territory length. To account for inter-experiment variability, mean intensity values for each condition were normalized to the average of the control condition within the corresponding batch, which was set to 100%

To quantify the percentage of c-Fos+ cells relative to Hoechst in Figure S4, images were thresholded in Fiji using a fixed automatic threshold method, applied consistently across images. Binary masks were processed with the "Fill Holes" function, and particles were counted using "Analyze Particles" with a size exclusion of 0.2–infinity, to exclude non-specific noise. The percentage of c-Fos+ cells was calculated as the ratio between c-Fos+ and Hoechst+ counts. In figure S5, the proportion of C-Fos positive cells were quantified using a custom-made plugin (https://github.com/orion-cirb/Cfos_2D_Syn_Dendrites). Briefly, for each image, the background of the GFP channel was subtracted, a threshold was applied and the image converted to a mask. This mask was applied to the C-FOS channel to measure the colocalization between Cthe C-FOS signal and the GFP signal.

SEMA6A and KI67 immunolabeling were acquired in Z stacks using Nikon Ti2 inverted microscope with CSU W1 (Yokogawa, Tokyo) spinning-disk scan head with 40x/1.3 Plan Fluor Oil objective using NIS Elements imaging software. SEMA- and KI67-staining thickness were measured and normalized to EGL thickness.

### Statistical Analysis

All imaging data were analyzed using GraphPad Prism for statistics. Values are presented as mean ± standard error of the mean (SEM). Normality was evaluated using the d’Agostino Pearson or Shapiro–Wilk normality test. When both conditions followed a normal distribution, the means between two conditions were compared using the unpaired Student’s t test. When data were normally distributed but their variances were different, Welch’s unpaired t-test was used. When the dataset did not follow a normal law, the means were compared using a Mann– Whitney nonparametric test. On column graphs, each dot represents one animal.

### Gene expression analysis RNA extraction

Cerebellum were dissected in cold HBSS. After removal of the meninges, the tissues were frozen in liquid nitrogen and stored at −80C°. Total RNA was extracted using the Qiagen RNeasy mini kit (Qiagen, Venlo, Netherlands, #74106) following tissue homogenization, according to manufacturer’s instructions. The quality of the RNA was evaluated using an Agilent RNA 6000 Pico kit (#5067-1513) and a 2100 Bioanalyzer Instrument.

### RNA sequencing

Bulk sequencing was performed at the ICGex - NGS platform at the Institut Curie using the Illumina paired-end 100nt approach and the stranded mRNA prep Ligation-Illumina protocol. Data management, quality control and primary analysis were performed by the Bioinformatics platform of the Institut Curie. Raw sequencing reads have been trimmed for adapter sequence and aligned on the Mouse mm10 reference genome (GENCODE vM22 annotation). For differential gene expression analysis, expressed raw counts were normalized using the TMM method from the edgeR package. Differential analysis was performed with the limma/voom R package. All raw p-values were then corrected for multiple testing using the Benjamini-Hochberg method.

### GO TERM analysis

Differentially expressed genes (DEGs) were separated into upregulated (logFC > 0) and downregulated (logFC < 0) gene sets. Genes with an adjusted p-value < 0.05 were included in downstream analysis. Gene Ontology (GO) enrichment analysis was performed using the cluster Profiler package (v4.20.0) independently for upregulated and downregulated gene sets. GO terms with an adjusted p-value < 0.05 and a q-value < 0.05 were considered significantly enriched.

The percentage of upregulated and downregulated genes was calculated for each cluster (=each differentiation stage) using for each the top 50 gene markers, identifying those that were amongst the DEGs identified by RNA sequencing, and estimating the proportion of total DEGs that corresponded to either up or downregulated. These percentages are visualized as a dot plot using the ggplot2 package (v.4.0.3), where dot size represents the percentage of genes in each regulation category.

### Spatial transcriptomics

Spatial transcriptomic analysis was performed using the Xenium In Situ platform (10x Genomics, Pleasanton, CA, USA) with the Xenium Prime 5K Mouse Pan Tissue and Pathways Panel supplemented with a custom add-on panel of 100 genes, according to the manufacturer’s protocol.

Briefly, 10-µm-thick fresh-frozen coronal cerebellar sections of P4 C57Bl/6J mice were mounted on Xenium slides and processed following the Xenium In Situ Gene Expression protocol. Fixation, permeabilization, probe hybridization, ligation, signal detection, cell segmentation, and transcript counting were performed using the Xenium platform and Xenium Analyzer. The spatial trancriptomic experiment using the Xenium technology was performed with the PHENOMIN - iCS facility, Illkirch, France.

### Analysis of the data

Data analysis was conducted in R (v4.5.2) using the Seurat package (v5.5.1). Gene expression data from P4 cerebellum were merged and filtered using the following criteria: nCount > 20 and nCount < 98th quantile. After quality filtering, log-normalization, scaling, dimensionality reduction, and clustering were performed using Seurat functions. Parameters for unsupervised clustering analysis were chosen using the clustree package (v0.5.1). To remove non-granule cell populations, module scores were calculated with AddModuleScore using curated marker gene sets representing Purkinje cells (*Ptf1a, Foxp2, Rora*), inhibitory interneurons (*Pax2, Gsx1, Ascl1*), Bergmann glia (*Sox2, Zeb2, Etv4, Sox9*), unipolar brush cells (*Eomes, Lmx1a*) and Golgi cells (*Lgi2, Grm2*). Cells with positive enrichment for any non-granule cell module score were excluded. The filtered granule cell lineage dataset was then re-normalized, scaled, dimensionally reduced and clustered. Cluster-specific marker genes were identified using Seurat’s FindAllMarkers function. GO enrichment analysis was performed on the DEGs using the clusterProfiler package. Marker genes with an adjusted p-value < 0.05 were retained for the analysis.

---

**Fig. S1.**
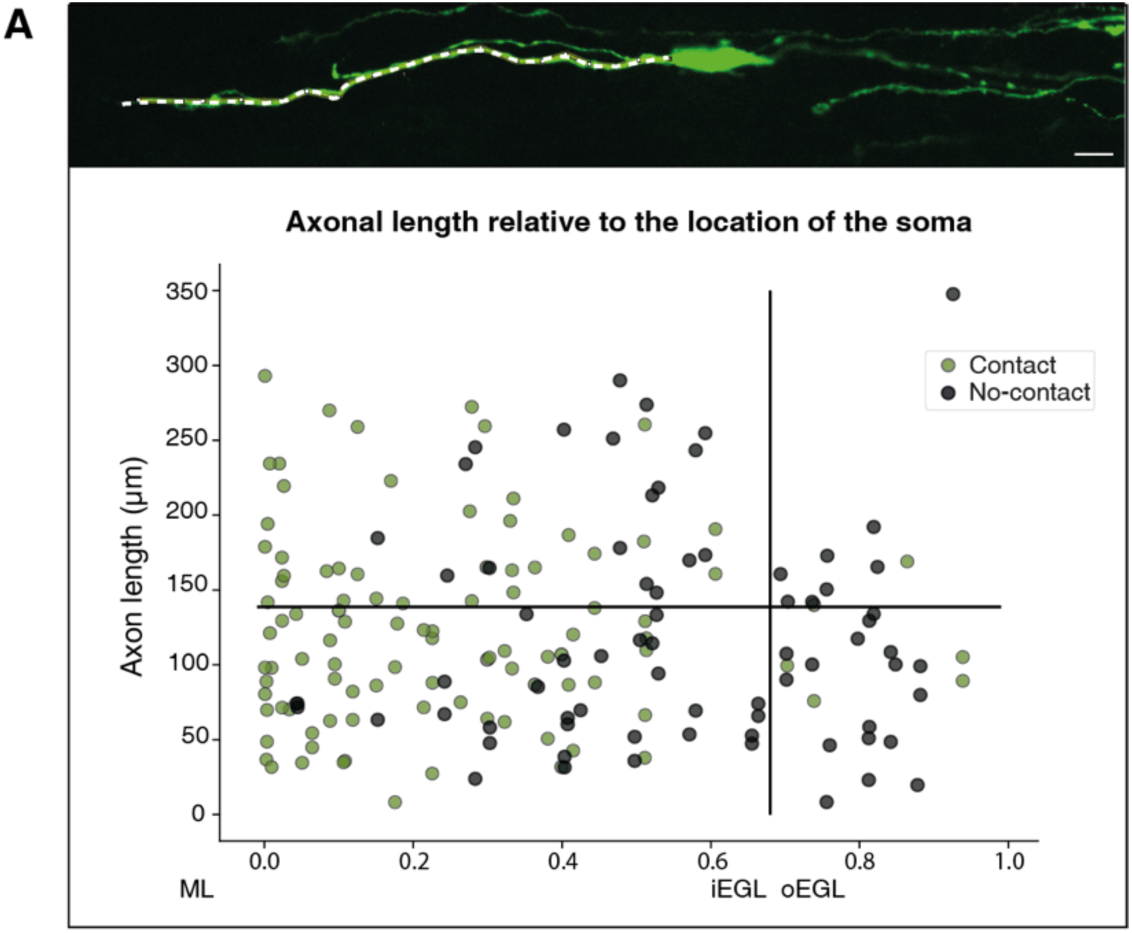
Axonal length is not correlated with the ability of differentiating Granule cells to contact Purkinje cells. (Top) Venus-positive granule cells are visualized using anti-GFP immunostaining. Axonal length is measured starting from the soma (white dashed line). Scale bar= 10 µm. (Bottom) Graph showing the axonal length for each labeled GC according to the distance of its soma relative to the ML border. GCs making contact are represented in green while GCs that don’t make any contact with PCs are represented in black. N = 5 independent experiments.

**Fig. S2.**
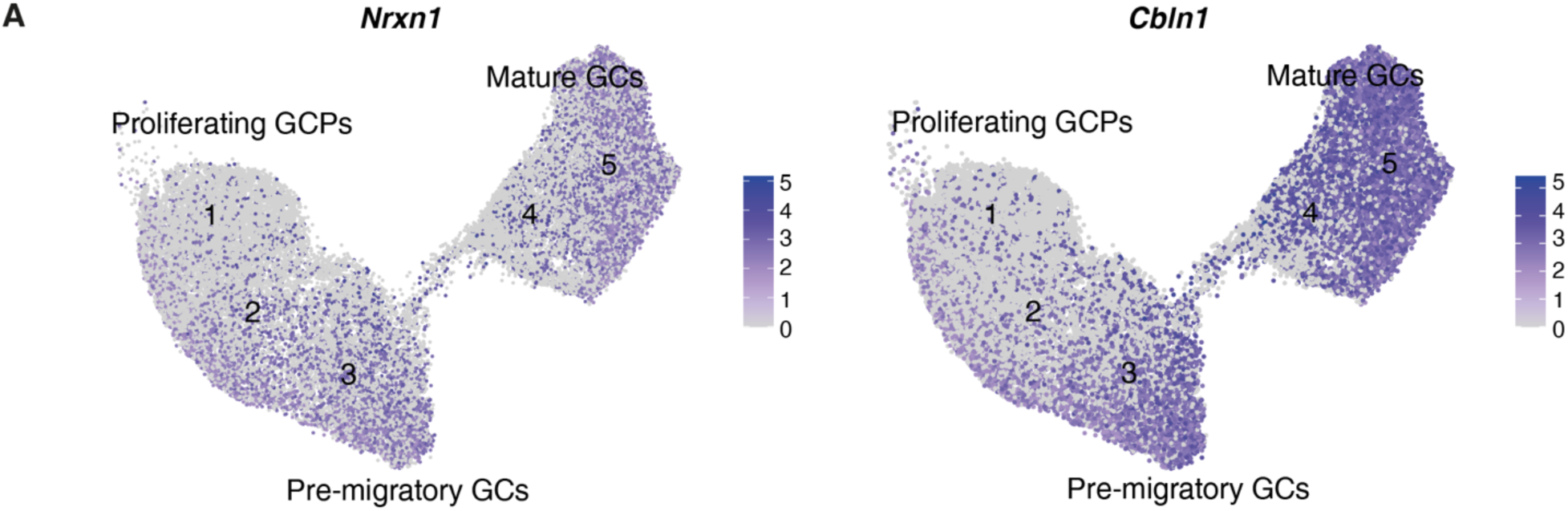
*Nrxn1* and *Cbln1* are expressed in pre-migratory GCs. Data from the spatial transcriptomic experiment show the *Nrxn1* and *Cbln1* expression patterns in the GC lineage in P4 mice. Clusters are annotated as in Figure 2.

**Fig. S3.**
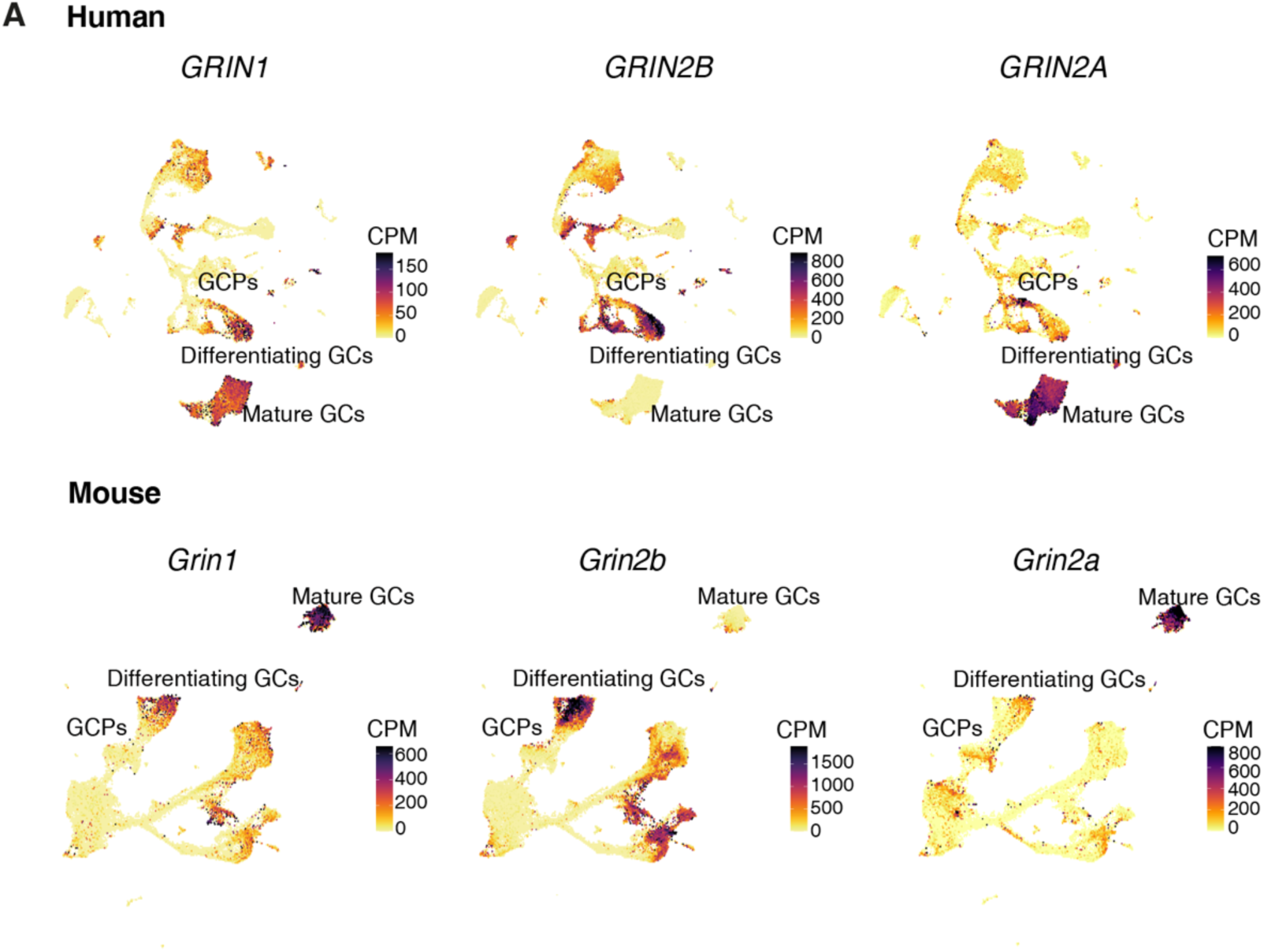
Gene expression of NMDA receptor subunits during human and mouse cerebellar development. (A) UMAP representation of gene expression patterns in the developing cerebellum of human and mice (single-nuclei transcriptomic data and representation from https://apps.kaessmannlab.org/sc-cerebellum-transcriptome/, (*46*))

**Fig. S4.**
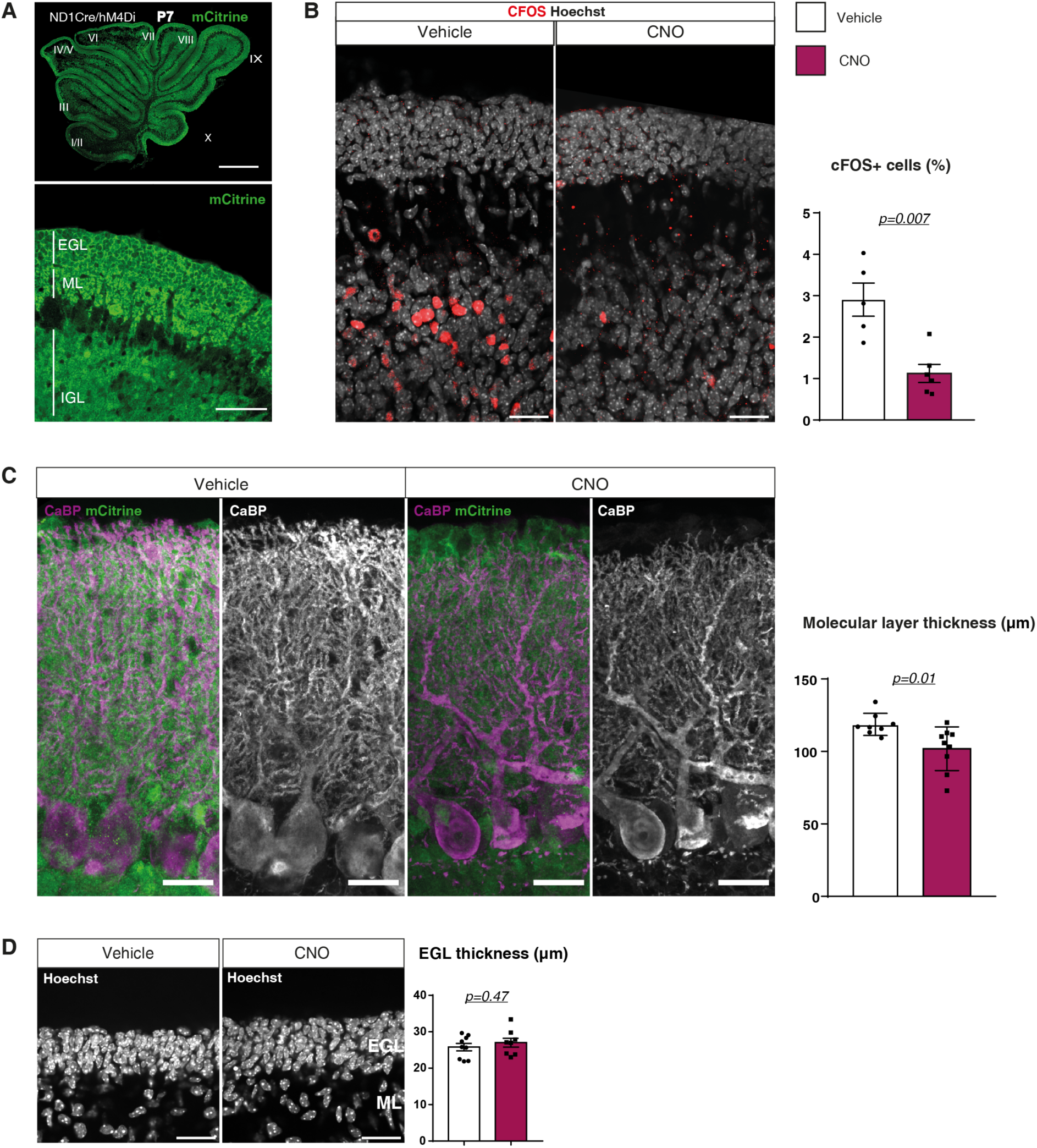
Characterization of the Neurod1-Cre;hM4Di mouse model. **(A)** Immunolabeling using an anti-GFP antibody (green) shows the expression of the reporter mCitrine in a parasagittal section of cerebellar vermis of a P7 NeuroD1Cre/hM4Di heterozygous mouse. Higher magnification in lobule IX (bottom) shows mCitrine in immature granule cells in the EGL, in mature GCs in the IGL, and in the ML that contains the parallel fibers. Scale bars: Top = 500 μm; bottom = 50 μm. **(B)** The percentage of C-Fos positive cells in the IGL of the cerebellar cortex was quantified after immunolabeling for C-Fos (red) and cell nuclei (Hoechst, gray) on P7 parasagittal cerebellar slices from NeuroD1Cre/hM4Di mice 1h30 after a single injection of CNO or vehicle (C-Fos cells -vehicle: mean of percentage 2.91 ± SEM 0.40, n=5 mice vs CNO: 1.13 ± SEM 0.22, n=6 mice; N = 2 independent experiments, Welch’s unpaired t-test p = 0.0072). **(C)** Anti-GFP immunolabeling shows expression of the reporter mCitrine in the granule cell layer and PFs in cerebellar sections of P14 NeuroD1Cre/hM4Di heterozygotes chronically treated with CNO or vehicle between P7 and P14. The dendritic tree and somata of PCs are immunolabeled using anti-CaBP (magenta) to enable measurement of the thickness of the ML (mean± SEM: vehicle= 118.7 ± 2.69 µm, n=8 mice vs CNO= 101.9 ±5.02 µm, n=9 mice; N = 2-3 independent experiments; p = 0.012 Welch’s unpaired t-test). Scale bars = 25 µm **(D)** EGL thickness was measured in vermal cerebellar sections of P11 vehicle or CNO injected Neurod1-Cre;hM4Di heterozygous mice, after stained using Hoechst to visualize the nuclei (mean ±SEM : vehicle=25.81 ±1.02 µm, n=9 mice vs CNO=27 ± 1.23 µm, n=8; N = 4 independent experiments, Welch’s unpaired t-test). Scale bars = 25 μm.

**Fig. S5.**
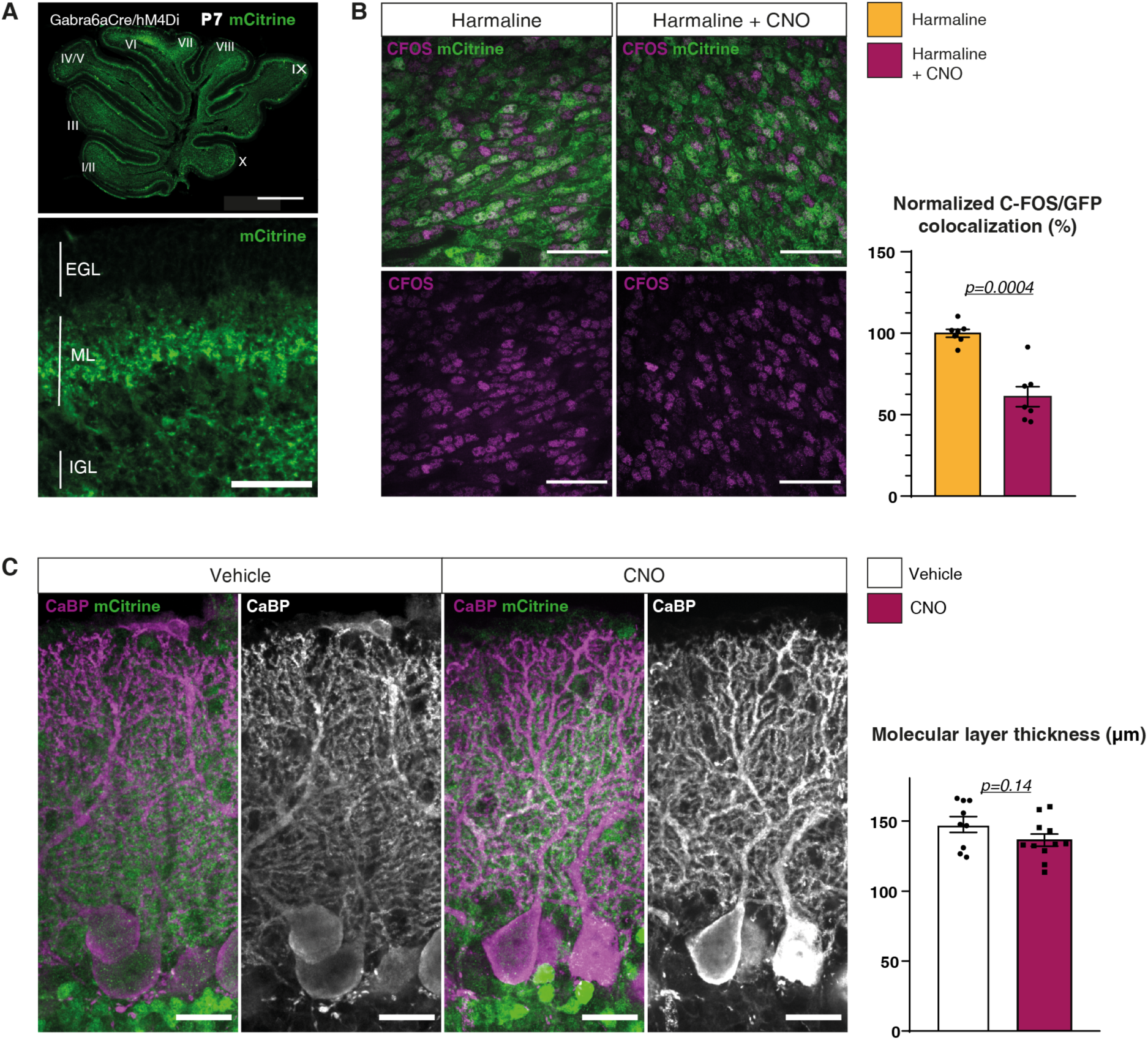
Characterization of the Gabra6-Cre;hM4Di mouse model. **(A)** Immunolabeling using an anti-GFP antibody (green) shows the expression of the reporter mCitrine in a parasagittal section of cerebellar vermis of a P7 Gabra6Cre/hM4Di heterozygous mouse. Higher magnification in lobule IX shows mCitrine presence in mature GCs in the IGL, and in the ML that contains the parallel fibers. Scales bars: Top= 500 μm; bottom= 50 μm. **(B)** Because of the low expression of the c-Fos protein in mature GCs, response to CNO in hM4Di expressing mature GCs from P12 mice was assessed by first injecting harmaline to increase activity in the olivocerebellar circuit (*55*) followed by CNO (or vehicle) administration 1h later. The percentage of c-Fos (magenta) positive cells in the mCitrine positive GC population was quantified (Normalized percentage±SEM: vehicle=100 ± 2.41, n=7 mice vs CNO=61.08 ±6.15, n=7 mice; N = 2 independent experiments, p = 0.0004 Welch’s unpaired t-test) **(C)** Anti-GFP immunolabeling shows expression of the reporter mCitrine in the granule cell layer and PF in cerebellar sections from P14 Gabra6Cre/hM4Di heterozygotes. The dendritic tree and somata of PCs are immunolabeled using anti-CaBP (magenta) to enable measurement of the thickness of the ML (mean± SEM: vehicle= 147.5 ± 5.61 µm, n=9 mice vs CNO=136.4 ±4.38 µm, n=11 mice; N = 3 independent experiments, p=0.14 Welch’s unpaired t-test).

